# Emotion suppression failures are associated with local increases in sleep-like activity

**DOI:** 10.1101/2020.08.04.235978

**Authors:** Giulia Avvenuti, Davide Bertelloni, Giada Lettieri, Emiliano Ricciardi, Luca Cecchetti, Pietro Pietrini, Giulio Bernardi

**Affiliations:** MoMilab, IMT School for Advanced Studies Lucca, Lucca, Italy; University of Pisa, Pisa, Italy

**Keywords:** EEG, emotion regulation, behavior, frontal cortex, sleep

## Abstract

Emotion self-regulation relies both on cognitive and behavioral strategies implemented to modulate the subjective experience and/or the behavioral expression of a given emotion. While it is known that a network encompassing fronto-cingulate and parietal brain areas is engaged during successful emotion regulation, the functional mechanisms underlying failures in emotion suppression are still unclear. In order to investigate this issue, we analyzed video and high-density EEG recordings of nineteen healthy adult subjects during an emotion suppression (ES) and a free expression (FE) task performed on two consecutive days. Changes in facial expression during ES, but not FE, were preceded by local increases in sleep-like activity (1-4Hz) in in brain areas responsible for emotional suppression, including bilateral anterior insula and anterior cingulate cortex, and in right middle/inferior frontal gyrus (p<0.05, corrected). Moreover, shorter sleep duration the night prior to the ES experiment correlated with the number of behavioral errors (p=0.01) and tended to be associated with higher frontal sleep-like activity during emotion suppression failures (p=0.05). These results indicate that local sleep-like activity may represent the cause of emotion suppression failures in humans, and may offer a functional explanation for previous observations linking lack of sleep, changes in frontal activity and emotional dysregulation.

## Introduction

Emotions are an essential aspect of the psychological life of human beings. In fact, they greatly affect our physiological, cognitive and behavioral responses to internal and external stimuli (Keltner and Kring, 1998; Cacioppo et al., 2000; Sapolsky, 2007; Koole, 2009). Importantly, emotions may occasionally lead to inappropriate or exaggerated reactions that could have negative consequences on our social life. Therefore, in dealing with emotions, people frequently engage in covert or overt forms of self-regulation in order to preserve a flexible and goal-oriented behavior (Koole, 2009; Kelley et al., 2015).

In general, self-regulation involves a balance between the strength of an impulse, its related reward, and the individuals’ ability to resist to the impulse and to modify their behavior in accordance to relevant personal goals (Carver and Scheier, 1998; Gross and Thompson, 2007). When applied to emotions, self-regulation typically implies adjusting their type, intensity, duration and expression (Gross, 1998). Various schemata, ranging from attention allocation to cognitive reappraisal and expressive suppression (Gross, 1998; Webb et al., 2012) can be used to modify an individual’s reaction to emotional states. For instance, when applying expressive suppression, a response-based modulation, individuals voluntarily refrain from overtly showing their emotional state, which is kept hidden to an external observer. Previous work investigated the neural correlates of voluntary emotion suppression by characterizing changes in brain activity during the presentation of emotion-inducing stimuli. The obtained results revealed that successful emotion suppression is associated with the activation of a broad fronto-parieto-insular network, including bilateral supplementary motor area (SMA), preSMA, anterior midcingulate cortex, anterior insula, inferior frontal gyrus, lateral orbitofrontal cortex, posterior middle frontal gyrus, dorsal temporo-parietal junction and the left posterior middle temporal gyrus (Frank et al., 2014; Kohn et al., 2014; Langner et al., 2018). Yet, what may cause a failure of this emotion-regulation system and the consequent generation of undesired behavioral responses still remains largely unclear.

Interestingly, sleep loss due to restriction or deprivation is known to significantly impair the ability to regulate emotional responses and affective states (Krause et al., 2017; Ben Simon et al., 2020a, 2020b), and these changes have been suggested to depend on an altered top-down control of the medial frontal cortex on limbic structures (Yoo et al., 2007). However, the actual functional cause of this frontal impairment and whether it may also explain emotion regulation failures observed in (apparently) rested wakefulness is unknown.

Recent evidence indicates that local, temporary intrusions of sleep-like brain activity may represent a common cause of behavioral errors when involving task-related brain regions (Hung et al., 2013; Bernardi et al., 2015; Nir et al., 2017; Slater et al., 2017; Quercia et al., 2018; Petit et al., 2019). Such local sleep-like episodes reflect spatially and temporally circumscribed reductions in neuronal firing (*off-periods)* and are associated with the appearance, in the electroencephalographic (EEG) signal, of delta-theta waves (≤8 Hz) similar to those of actual sleep (Vyazovskiy et al., 2011). Given that they increase in number and extension as a function of time spent awake and that such changes are reverted by a night of sleep, local sleep-like episodes have been suggested to represent a signature of brain functional ‘fatigue’ and a direct cause of behavioral impairment following sleep loss (Andrillon et al., 2019; D’Ambrosio et al., 2019). Of note, during wakefulness, the number of sleep-like episodes does not increase homogeneously over the cortical mantle. In fact, frontal areas typically display the strongest increases in low-frequency activity relative to other brain regions, suggesting a particular vulnerability to functional fatigue (Finelli et al., 2000; Strijkstra et al., 2003).

In light of the above considerations, here we hypothesized that local sleep-like episodes occurring in areas of the emotion regulation network, and especially within frontal brain regions, could represent a potential cause of emotion suppression failures. In particular, we predicted that a shorter sleep time or a reduced sleep quality could lead to a higher incidence of frontal sleep-like episodes the following morning, which would in turn result in higher probability of emotion suppression failures.

## Methods

### Subjects

Nineteen healthy adults (age range = 21-31 years, mean ± SD = 26.3 ± 2.9 years, 10 females, all right-handed) were included in the study. All participants underwent a preliminary interview to exclude any clinical, neurological or psychiatric conditions potentially affecting brain function and behavior. Additional exclusion criteria comprised excessive daytime sleepiness (Epworth Sleepiness Scale score >10; Johns, 1991) and extreme chronotypes (Morningness-Eveningness Questionnaire score >70 or <30; Horne and Ostberg, 1976). Participants were asked to maintain a regular sleep-wake schedule for at least one week before each experiment. Compliance was verified by wrist-worn actigraphy (MotionWatch 8, CamTech). The study was conducted under a protocol defined in accordance with the ethical standards of the 2013 Declaration of Helsinki and approved by the Local Ethical Committee. Written informed consent was obtained from all participants.

### Experimental Procedures

Data analyzed in the present work was collected as part of a larger study aimed at investigating the neural and behavioral consequences of extended task practice (*unpublished*). A general overview of the whole experimental protocol is provided herein.

All participants completed a practice session and two experimental visits in which EEG (EGI, Eugene, OR, USA; 64 electrodes, 500 Hz sampling rate) activity and behavioral data were recorded. In order to minimize inter-individual differences in wake-sleep rhythms and work-related fatigue, all sessions were performed with a pre-defined, fixed schedule. In particular, the practice session was performed on Friday morning from 9:30 AM to 11:30 AM, while the two experimental visits took place on the next Monday and Tuesday, from 8:30 AM to 1:00 PM.

During the practice session, participants completed five 5-min trials of a motor-response inhibition task (Garavan et al., 1999, 2002; Roche et al., 2005; Chuah et al., 2006; Bernardi et al., 2015; 300 stimuli, 10% lures). Data obtained from this procedure was used to calibrate the difficulty of the same task presented during the two subsequent experimental sessions, as described in previous work (Chuah et al., 2006).

The two experimental visits required the participants to complete partially different versions of the same tasks. Each experiment began with a ∼15-min long test block (baseline; BL) including two 2-min resting-state EEG recordings with eyes open (4 min in total) and two trials of the response inhibition task. Moreover, subjective vigilance, sleepiness, mood, perceived effort and motivation were assessed using 10-point Likert scales. This initial assessment was followed by three ∼45-min long task sessions (TS1-3), each one followed by a ∼15-min long test block identical to the first one (Figure 1). In one of the two experimental visits the task sessions included three computerized tasks requiring high levels of impulse control, decision making and conflict resolution, i.e., an emotion suppression task (Baumeister et al., 1998; Dang, 2018), a false response task (Bernardi et al., 2015), and a classical Stroop task (Stroop, 1935). A detailed description of the emotion suppression task, which was the focus of the present study, is provided below. In the other experimental visit, a modified version of the same tasks was presented, which required no exertion of self-control (e.g., Stroop task with consistent word-ink color). Finally, before the last task block a caffeinated or decaffeinated beverage was provided to participants. The order of the two experimental visits and the administration of caffeinated vs. decaffeinated beverages were randomized across participants. All computerized tasks were implemented using E-Prime 2 (Psychology Software Tools, Pittsburgh, PA).

**Figure 1.**
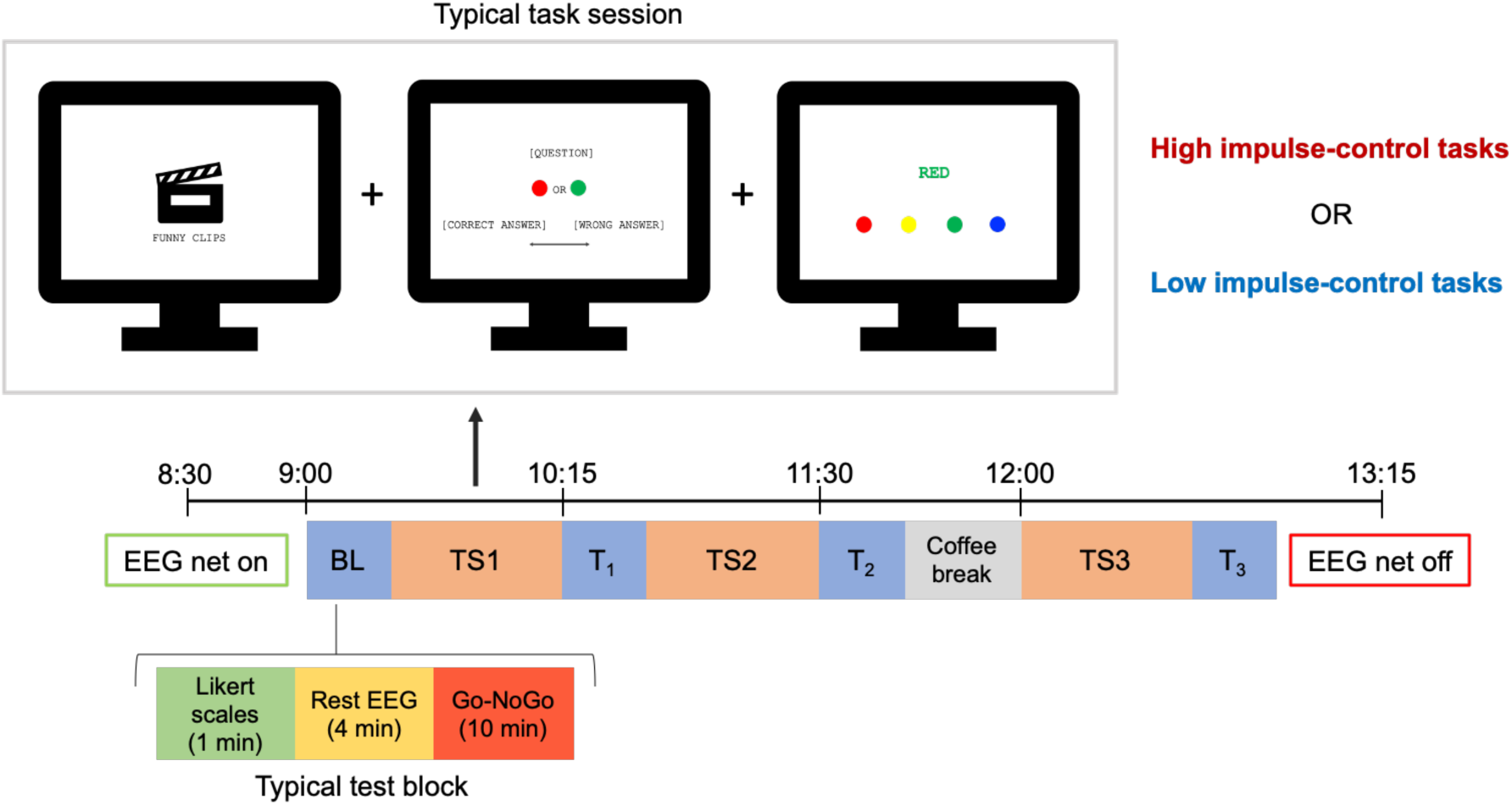
Experimental paradigm. The baseline assessment (BL) and each test block (T_1_-T_3_) included Likert scales, resting-state EEG recordings and the completion of two trials of a response inhibition task (Go-NoGo). In one of the two experimental visits (high impulse-control effort condition), the subjects completed an emotion suppression task, a false response task and a Stroop task. During the other experimental visit (low impulse-control effort condition), the participants completed a simplified version of the same tasks requiring no exertion of self-control.

### Emotion suppression/expression task

In one experimental visit, participants completed three instances (one for each task session) of an emotion suppression task in which they were explicitly requested to suppress their facial reactions while watching amusing video-clips (‘emotion suppression’ condition, ES). In the task sessions of the other experimental visit, volunteers watched similar video-clips, but they were left free to express their emotional responses (‘free expression’ condition, FE). In both conditions, the participants’ faces were video-recorded using a camera positioned above the PC-monitor and synchronized with the software used to record the EEG signals (NetStation 5.3, EGI). Of note, for each task session, the video-clips were presented in two trials in which they were alternated with 4 s of black frames, for an average total duration of ∼6 min (5.9 ± 0.3 min).

In order to present different contents to participants during each task session, a total of 261 clips depicting people and/or animals in amusing situations were downloaded from the internet. An initial validation procedure was performed in an independent sample of 12 subjects (age range = 23-36 years, mean ± SD = 29.6 ± 4 years, 8 females) to ensure a balanced distribution of the emotional contents across sessions. Specifically, the volunteers were asked to rate the set of amusing clips and an additional set of 20 clips showing simple actions/activities with no obvious emotional content. The clips were presented in random order. After each video, the subjects answered the question *“How difficult would it be for you to keep a neutral expression while watching this clip?”* using a rating scale ranging from 1 (*“Not difficult at all”*) to 5 (*“Very difficult”*). As expected, the 261 amusing clips were associated in each subject with significantly higher ratings with respect to the 20 neutral clips (p < 0.001). For each subject, the ratings given to the amusing video-clips were re-scaled by computing the ratio with respect to the average rating of the 20 neutral clips, which were considered as an individual baseline. Then, the group-average ratios were computed for each clip, and a randomization procedure was used to assign the clips to each task trial, thus ensuring a similar distribution of ratings and a similar total duration of the video stimuli (Table 1).

**Table 1.** Mean and standard deviation (SD) of video ratings (ratio with respect to mean score of neutral videos) for each task trial, presented during the ‘emotion suppression’ condition (ES) and the ‘free expression’ condition (FE). No statistically significant differences were found across experimental conditions.

|  | ES | FE |  |  |
| --- | --- | --- | --- | --- |
| | mean ± SD | mean ± SD | z-score | p-value |
| **TS1-1** | 1.85 ± 0.23 | 1.90 ± 0.20 | 0.393 | 0.705 |
| **TS1-2** | 1.94 ± 0.29 | 1.87 ± 0.25 | 0.608 | 0.551 |
| **TS2-1** | 1.88 ± 0.27 | 1.97 ± 0.17 | 0.718 | 0.488 |
| **TS2-2** | 1.88 ± 0.25 | 1.82 ± 0.28 | 0.464 | 0.661 |
| **TS3-1** | 1.92 ± 0.16 | 1.91 ± 0.24 | 0.083 | 0.946 |
| **TS3-2** | 1.88 ± 0.18 | 1.87 ± 0.26 | 0.103 | 0.924 |

### Scoring of video recordings

In order to identify and quantify the occurrence of facial expression changes in each task trial, video recordings of each participant were visually inspected and scored by one of the authors (D.B.). The scoring procedure was performed in two steps. First, the operator watched each video using a custom-made program written in Psychtoolbox v3.0.1 (Kleiner et al., 2007) for MATLAB (v9.7; Natick, Massachusetts: The MathWorks Inc.). Each time a change in facial expression was identified, the scorer pressed a button and the corresponding time-point (in milliseconds) was stored. In the second step, another custom-made MATLAB program was used to re-inspect, frame-by-frame, the time-period around the tagged time-point (± 2 s) and to select the video-frame that immediately preceded the change in facial expression. In this step, each event was also accurately re-evaluated and all cases in which a clear change in facial expression was not confirmed were marked for rejection. For this whole procedure, the scorer remained blind to the experimental condition of each video. Finally, tagged events that occurred after 1 second from the end of a clip and before the beginning of the subsequent clip (i.e., during inter-stimulus black frames) were also automatically discarded.

### EMG data analysis

A facial electromyographic (EMG) signal was derived from two EEG electrodes located below the two eyes, on the cheeks, approximately above the zygomatic muscles. The continuous signal of these two channels was referenced on homolateral, pre-auricular electrodes, and band-pass filtered between 30 and 200 Hz. A notch filter at 50 Hz was also applied.

Variations in facial EMG activity were evaluated to confirm the expected association between tagged events and changes in facial expression. Moreover, given that the estimation of expressive changes based on visual inspection of the video-clips may be inaccurate with respect to the actual beginning of muscular activity, the EMG signal was also inspected to determine the specific onset of each tagged event (Fiacconi and Owen, 2015). In particular, for the EMG inspection procedure, the root mean square (RMS) of the filtered signals was computed using a moving-window approach (1-s length; 1 time-point steps). Then, 8-s long data segments, including 4 s before and 4 s after the manually tagged onset of changes in facial expression were extracted. Each individual event was visually inspected using a custom-made MATLAB function and the onset of the increase in EMG activity was marked. Cases for which a clear increase in EMG activity were not identified were excluded from further analyses. Finally, for all the retained episodes, 8-s long data segments centered on the EMG-activation onset were extracted and the total signal power (30-200 Hz) was computed in 2-s epochs using the Welch’s method (*pwelch* function, MATLAB signal processing toolbox; Hamming windows, 8 sections, 50% overlap). The mean power in the 4 s after onset-time were compared to the mean signal power in the 4-s epoch preceding the same time-point at group level (i.e., after within-subject averaging across episodes). The ratios between the two data segments (post/pre) were also compared across experimental conditions (ES, FE) to investigate potential differences.

### EEG data analysis

Continuous EEG recordings performed during the emotion suppression/expression task were band-pass filtered between 0.5 and 45 Hz. All EEG traces were visually inspected to identify and mark bad channels containing clear artifactual activity using NetStation 5.3 (EGI). Then, an Independent Component Analysis (ICA) was performed in EEGLAB (Delorme and Makeig, 2004) to remove signal components reflecting ocular, muscular and electrocardiograph artifacts. Rejected bad channels were subsequently interpolated using spherical splines. After preprocessing, all EEG traces were re-referenced to average reference and 4-s-long data epochs immediately preceding the EMG-activation onset (t = 0 s) of changes in facial expression were extracted. Finally, for each subject, the signal power in delta (1-4 Hz) and theta (4-8 Hz) frequency-bands was computed for each epoch as described for the EMG signal and averaged across episodes of interest. Paired comparisons between experimental conditions were then performed at group level.

Of note, given that potential power differences between experimental conditions could emerge due to differences in task demands (i.e., emotion suppression vs. free expression of emotions), further analyses were planned to exclude differences in overall brain activity across ES and FE or a direct correlation between low-frequency brain activity and the task-related effort required to suppress emotional expressions. In particular, for the first analysis, the mean (or median) signal power computed across non-overlapping 2-s epochs covering each task-related EEG recording was compared across experimental conditions. The second analysis was performed only on data of the ES condition. Here, for each subject, we calculated the Spearman’s correlation coefficients between mean signal power values computed in 2-s epochs during the presentation of each video-clip and the corresponding ratings expressing the effort needed to suppress emotional responses. All video-clips that were associated with a change in facial expression were excluded from this computation. A one-sample test based on z-scored transformed correlation values was performed to identify potential group-level effects.

### Source modeling of EEG data

The signal of pre-processed EEG-epochs corresponding to the 4 s immediately preceding the onset of changes in facial expression were source modeled using *Brainstorm* (Tadel et al., 2011). Specifically, the conductive head volume was modeled using a three layers symmetric boundary element method (OpenMEEG BEM, Kybic et al., 2005; Gramfort et al., 2010) and the default ICBM152 anatomical template. A standard set of electrode positions (GSN HydroCel 64) was used to construct the forward model. The source space was constrained to the cerebral cortex, which was modeled as a three-dimensional grid of 15,002 vertices. The inverse matrix was computed using the standardized low-resolution brain electromagnetic tomography (sLORETA) constraint with a regularization parameter equal to 10^−2^ λ. Finally, the signal power was computed for each vertex in source space using the Welch’s method (1-s Hamming windows, 50% overlap). Power maps in source space were then exported to MATLAB for planned statistical comparisons and obtained results were re-imported in *Brainstorm* for visualization.

### Data selection

In order to avoid potential confounding effects related to extended practice with partially different tasks in the two experimental conditions (ES, FE), as well as to the administration of caffeinated or decaffeinated beverages in the last hour of the two experiments, our analyses of behavioral and EEG/EMG data collected during the emotion suppression/expression task specifically focused on the first task session (i.e., TS1-1 and TS1-2). However, confirmatory analyses of relevant EEG-based comparisons were performed using all available task sessions, with and without application of a within-trial normalization. In particular, the normalization of EEG-signal power was performed by subtracting the median band-specific power computed across the whole corresponding EEG recording (2-s data segments; *pwelch* method with Hamming windows, 8 sections, 50% overlap).

### Statistics

Comparisons of EEG-signal power across experimental conditions were performed using paired t-tests and a permutation-based supra-threshold cluster correction, as described in previous work (Nichols and Holmes, 2002; Huber et al., 2006). In brief, the same statistical contrast was repeated after shuffling the labels of the two experimental conditions and the maximum size of significant electrode-clusters (p < 0.05) was saved in a frequency table. A minimum cluster-size threshold corresponding to the 95th percentile of the resulting distribution was applied to correct for multiple comparisons. For comparisons performed at scalp-level, the analysis was restricted to 51 ‘internal’ electrodes as described in previous work (Hung et al., 2013). In fact, more ‘external’ channels, located near the eyes, or on the temporal or neck muscles, are more likely to be affected by potential residual artifactual activity. Finally, permutation tests were used to assess the statistical significance of comparisons regarding behavioral and subjective variables and of correlations computed using the Spearman’s correlation coefficient. Of note, for all cases in which the number of possible data recombinations was greater than 10,000 this value was used to approximate the null distributions. In all other cases, the exact number of possible data recombinations was used. All statistical analyses were performed in MATLAB.

## Results

### Behavioral results

During the first task session (TS1), all subjects showed a lower number of facial expression changes in ES (2.6 ± 3.2) relative to FE (20.2 ± 10.7) condition (p < 0.001, |z| = 3.767; Figure 2A). Importantly, we found a negative correlation between actual sleep time in the night preceding the ES experiment and the absolute number of emotion suppression failures (p = 0.0126, r = −0.57; Figure 2B). A consistent but non-significant correlation was also found with subjective sleepiness (p = 0.119, r = −0.37), so that higher sleepiness was positively related to the number of suppression failures. Of note, self-reported mood showed instead no relationship with self-control failures (p = 0.306, r = −0.25).

**Figure 2.**
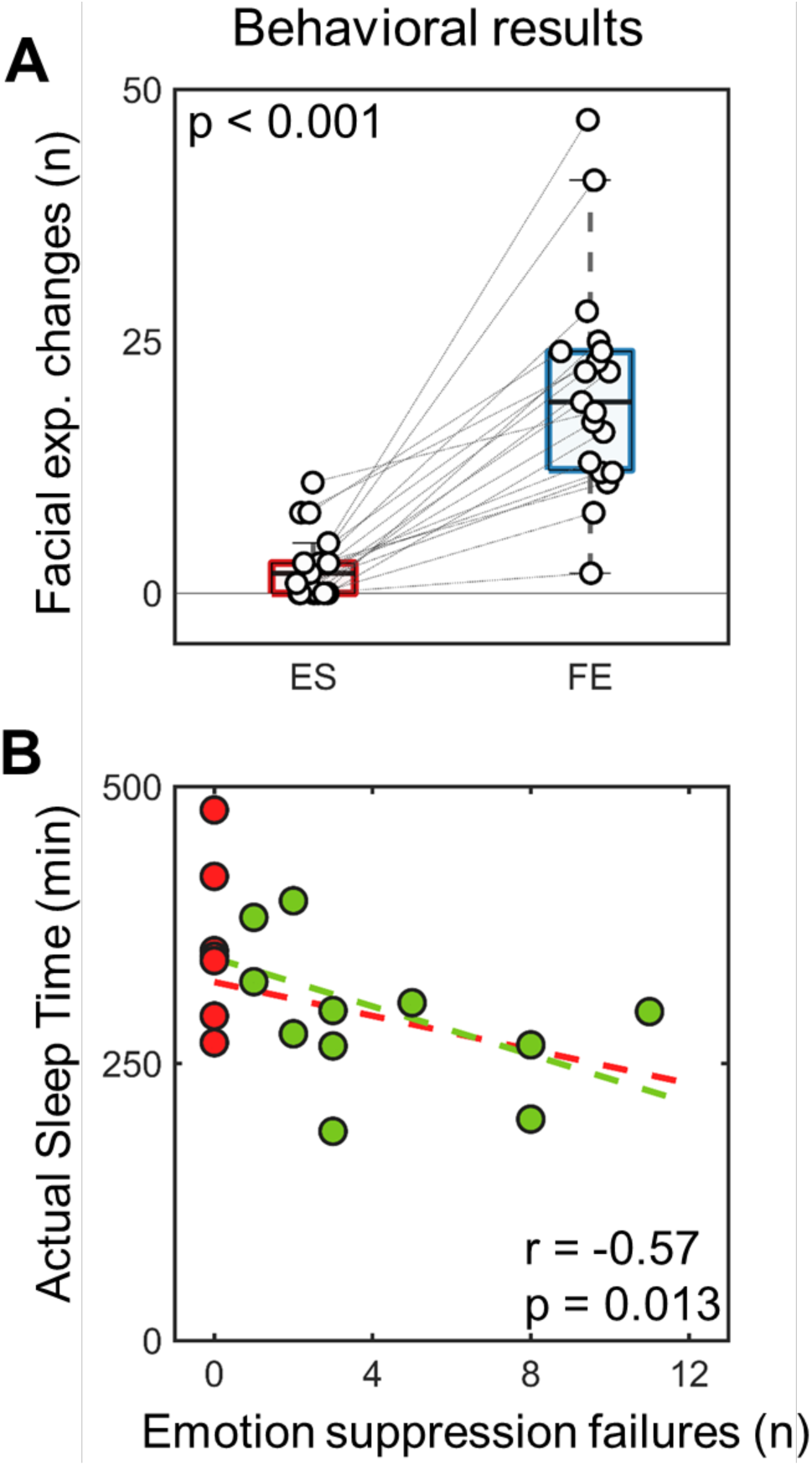
(A) Changes of number of facial expressions in the two experimental conditions (ES = emotion suppression; FE = free expression). (B) Relationship between number of emotion suppression failures and actual sleep time the night preceding the experimental session. This analysis was performed on 18 subjects due to missing actigraphic data in one participant (Spearman’s rho). The subjects who showed no changes in facial expression during TS1 are marked with red color. The remaining subjects are displayed using green color. Dashed lines represent the least-square fit for the distributions including all subjects (red) or only subjects who had at least one facial expression change (green).

Twelve participants (mean age = 26.7 ± 3.1, 8 males) had at least one emotion suppression failure in the ES condition and were thus included in further analyses. Specifically, these subjects had on average 4.1 ± 3.2 facial expression changes in the ES condition; a value still significantly lower than the one observed in the FE condition (22.9 ± 11.4; p < 0.001, |z| = 2.980). In the same subjects, the analysis of variations in EMG activity (30-200Hz) of the zygomatic muscle confirmed that marked facial expression changes were associated with significant activity increases in both experimental conditions (ES: p < 0.001, |z| = 1.682; FE: p < 0.001, |z| = 2.581; Figure S1). Relative EMG changes tended to be stronger in ES (p = 0.029, |z| = 1.842), although values also showed more variability across subjects in this particular condition. In fact, a visual re-inspection of video-recordings in ES confirmed a broad variability in the manifestation of suppression failures, with some subjects showing ‘explosive’ losses of control, and others only presenting minimal changes in facial expression. Instead, reactions tended to be more similar across subjects during FE.

### Sleep quality, vigilance and mood

Several control analyses were performed to exclude potential systematic differences between experimental conditions in the twelve examined participants. First, we investigated possible differences in time spent in bed and in actual sleep time between the nights that respectively preceded ES and FE experiments (of note, this analysis was performed on 11 subjects due to missing actigraphic data from one volunteer; age 28 yrs, male). We found no evidence of systematic differences (time spent in bed: p = 0.341, |z| = 1.004; actual sleep time: p = 0.492, |z| = 0.720; Figure S2A). Similarly, we found no systematic differences in sleepiness (p = 0.563, |z| = 0.904; Figure S2B), mood (p = 0.766, |z| = 0.632; Figure S2C) and motivation (p = 0.500, |z| = 1.134; Figure S2D). Of note, similar results were obtained when considering 18 subjects for actual sleep time (missing data in one subject) and all 19 subjects for the Likert scales (*data not shown*).

### Delta and theta EEG activity

We next evaluated whether increases in low-frequency (delta/theta) activity preceded the occurrence of emotion suppression failures (Figure 3). To this aim we compared the signal power computed in the four seconds preceding a change in facial expression across ES and FE conditions. We found that changes in facial expression were preceded by an increase in delta activity (1-4Hz) in frontal and left temporo-parietal electrodes (p < 0.05, corrected). These results were confirmed in additional analyses that included 15 subjects (mean age = 26.6 ± 2.8, 9 males) who had at least one emotion suppression failure across the three task sessions (TS1-TS3) of EF (p < 0.05, corrected; Figure S3). No significant differences in theta activity (4-8Hz) were found.

**Figure 3.**
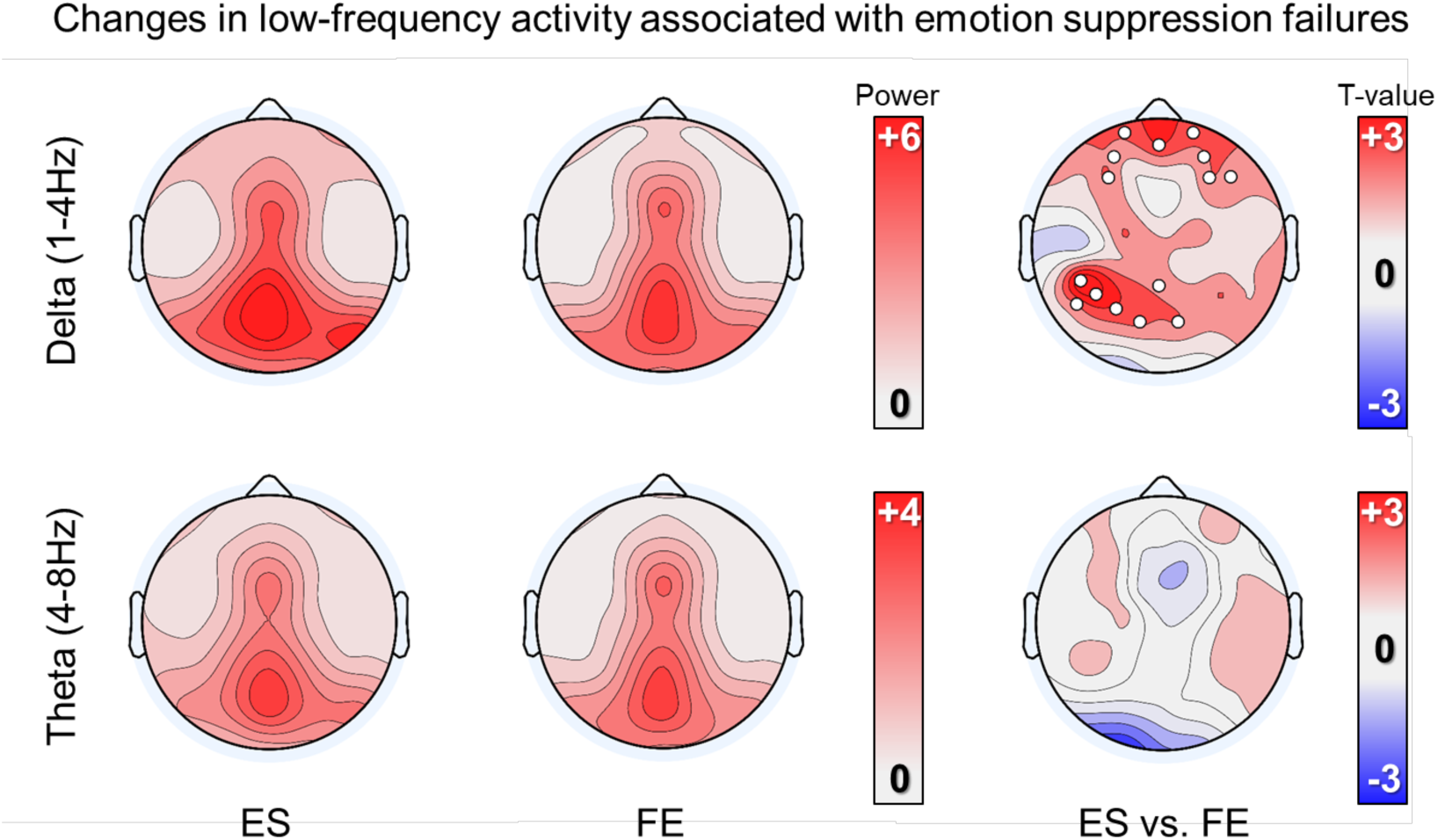
Low-frequency EEG activity associated with emotion suppression failures. Topographic plots on the left show absolute values of delta (top) and theta (bottom) power for the two experimental conditions (ES, FE) in the 4 s that preceded changes in facial expression. Topographic plots on the right show the statistical comparison between experimental conditions for the two frequency bands. White dots mark p < 0.05, cluster-based correction.

Given that changes in low-frequency activity may represent a signature of brain functional fatigue caused by insufficient sleep, we then explored the possible relationship between delta activity and actual sleep time the night preceding the ES experiment. We found that maximum delta power in frontal electrodes tended to be correlated with sleep time (p = 0.054, r = −0.60). Thus, a lower amount of sleep tended to be associated with higher levels of failure-related delta activity. Of note, a similar relationship was not found for temporo-parietal electrodes (p = 0.664, r = 0.15). Consistent but relatively weaker results were found using subjective sleepiness instead of actual sleep time (frontal: p = 0.066, r = 0.55; parietal: p = 0.198, r = 0.39).

### Possible confounding effects of task demands

Differences in delta power between experimental conditions could have emerged because of differences in task demands. Additional analyses were thus performed to investigate this possibility. First, we compared the mean/median delta power of the entire recordings of ES and FE to identify potential differences in overall task-related brain activity. This analysis revealed no significant differences between ES and FE either using the median (Figure 4A) or the mean (*data not shown*) power across epochs (p < 0.05, corrected). Next, we evaluated whether delta power was correlated with the effort required to suppress emotional responses in the EF conditions, as determined based on the scores provided by an independent sample of subjects (Table 1). No significant correlations were found between delta power and effort scores (p < 0.05, corrected; Figure S4), again indicating that differences in delta power do not reflect task-related demands in the ES condition.

**Figure 4.**
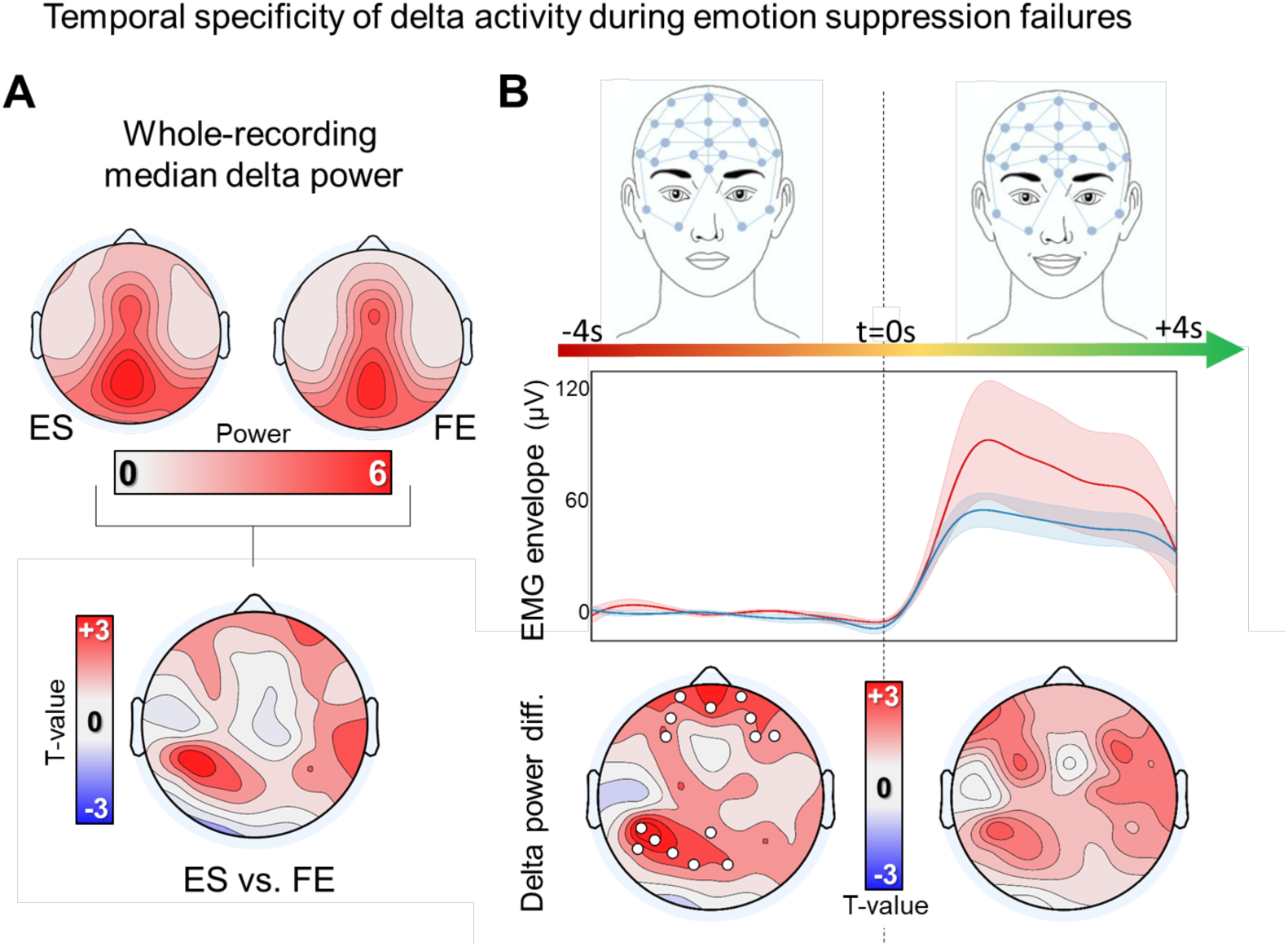
(A) Median task-related delta power computed for the entire EEG recordings. The topographic plot at the bottom shows the contrast between delta activity of ES and FE. (B) Contrast between experimental conditions in the 4 seconds before and in the 4 seconds after the change in facial expression (t = 0s). The central graph shows the variation of EMG activity in the same time window (envelope of rectified filtered EMG-signal). Values were normalized before averaging by subtracting the mean EMG activity computed in the 4 seconds before the onset of changes in facial expression (t = 0s). Shaded areas correspond to the standard error of the mean. For topographic plots, white dots mark p < 0.05, cluster-based correction.

### Temporal and frequency-band specificity

Additional analyses were performed to verify whether the observed association between changes in delta activity and emotion suppression failures were temporally and frequency-band specific. As described above, a temporal specificity is already suggested by the lack of significant differences between ES and FE for overall task-related delta power (Figure 4A; also see Figure S3). In addition, we found no significant differences between the two experimental conditions when comparing delta activity in the 4 s after the onset of changes in facial expression (Figure 4B; p < 0.05, corrected).

Next, we evaluated whether additional differences between ES and FE were present in other frequency bands (Figure 5). In line with previous results, we found no significant differences for alpha (8-12Hz), sigma (12-16Hz) and beta (18-30Hz) activity.

**Figure 5.**
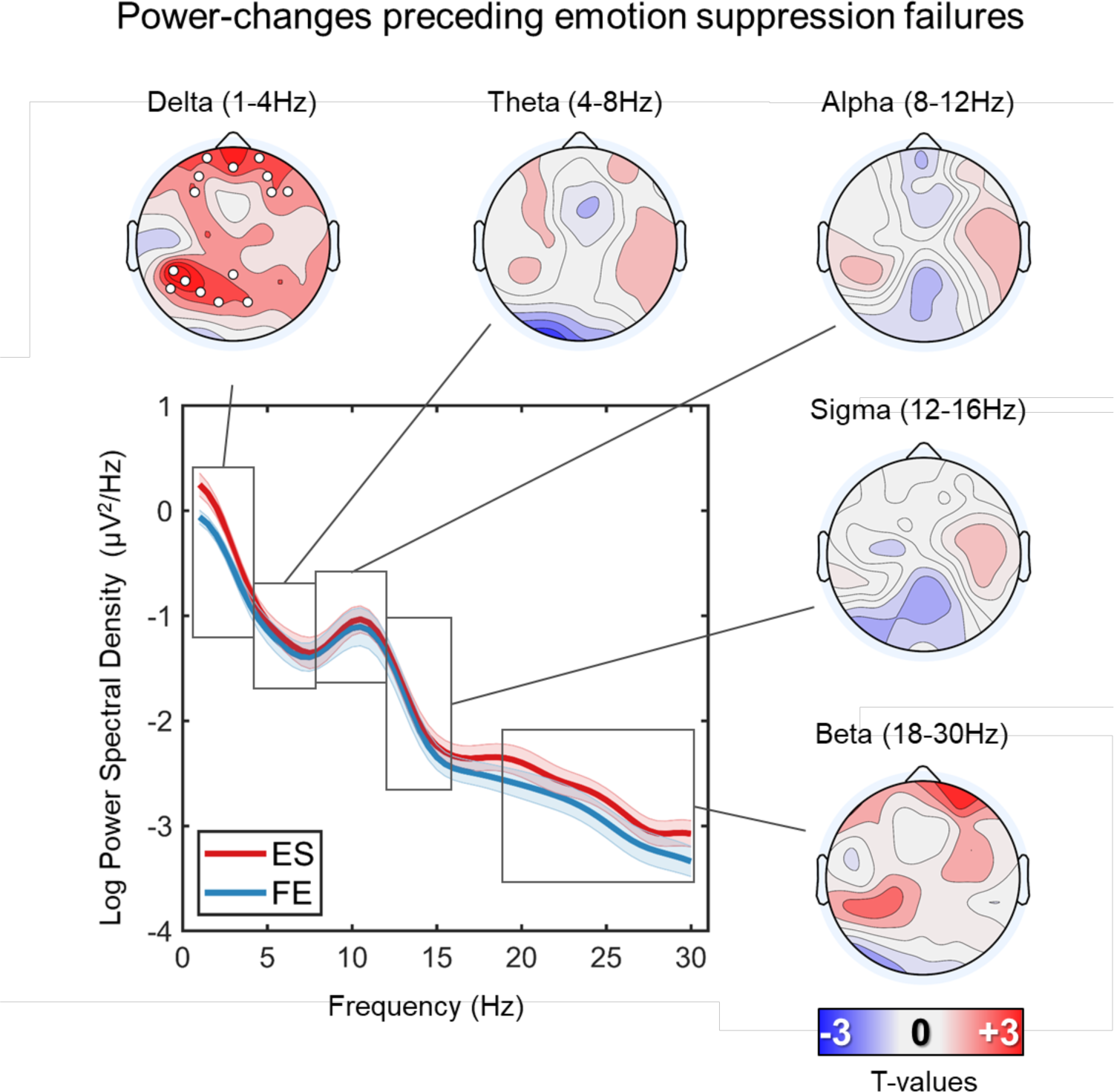
Differences in brain activity preceding the onset of changes of facial expression in the two experimental conditions (ES, FE). The central plot shows the power spectral density (PSD) computed across the electrodes presenting a significant difference in delta activity between ES and FE (white dots). Additional comparisons were performed for other typical frequency bands, including theta, alpha, sigma and beta. White dots mark p < 0.05, cluster-based correction.

### Source modeling analysis of delta activity

In order to identify the actual sources of changes in delta activity observed in the ES condition, the same analysis shown in Figure 3 was repeated after source reconstruction of the EEG signals (sLORETA; but very similar results were obtained using dynamical Statistical Parametric Mapping, dSPM). The obtained results are shown in Figure 6 (p < 0.05, corrected; cluster-forming threshold set to uncorrected p < 0.001; also see Figure S5). In particular, we found significant clusters in the bilateral anterior cingulate and medial frontal cortex, the anterior insula, the right precuneus and the right motor/premotor cortex, including the middle and inferior frontal gyri. Of note, most of these regions overlap with key areas previously described as part of the emotion regulation network (Langner et al., 2018).

**Figure 6.**
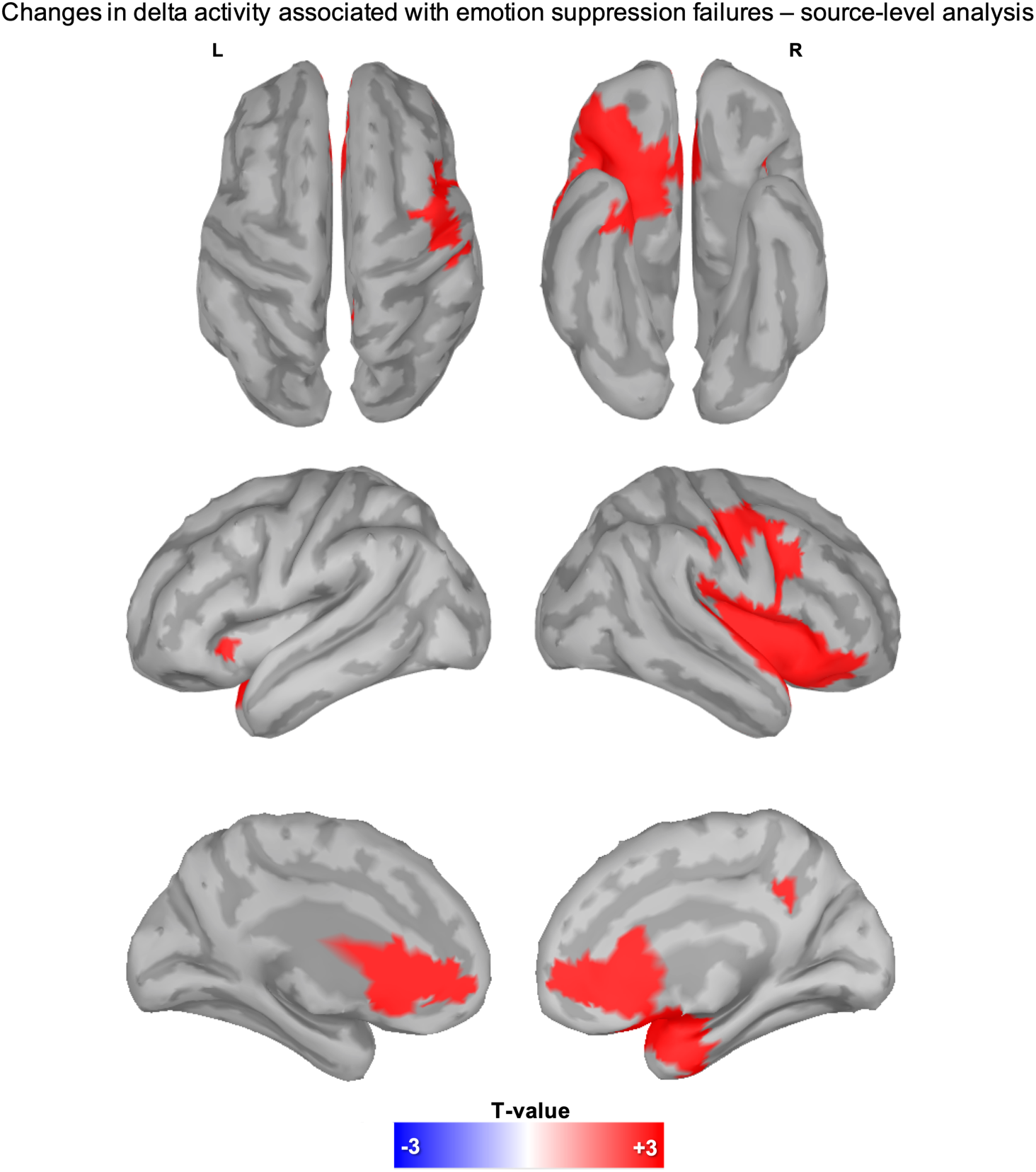
Delta EEG activity associated with emotion suppression failures. Significant differences (p < 0.001, cluster-based correction) in delta activity between ES and FE in the 4 seconds that preceded the onset of changes in facial expression.

## Discussion

Previous evidence indicates that frontal, parietal and limbic brain regions are activated during successful emotion suppression (Dörfel et al., 2014; Frank et al., 2014; Kohn et al., 2014; Langner et al., 2018). However, the neural correlates of emotion suppression failures were unknown. Here we demonstrated that self-control failures in an emotion suppression task are preceded by increases in low-frequency, sleep-like activity over frontal, insular and parietal regions. In addition, a positive correlation was found between actual sleep time the night before practice with the emotion suppression task and the absolute number of self-control failures, so that shorter sleep duration was associated with a poorer behavioral performance. These results indicate that intrusions of sleep-like activity in the brain network responsible for emotion regulation affect the efficacy of emotion suppression in healthy individuals. In this light, the occurrence of local, sleep-like episodes may offer a possible neurophysiological mechanism for previous observations regarding the effects of sleep loss on emotion regulation.

### Neural correlates of emotion suppression failures

Previous work showed that a neural network encompassing frontal, cingulate and parietal regions is recruited during emotion suppression tasks (Langner et al., 2018). However, to the best of our knowledge, no study investigated the neurophysiological events that could underlie failures in emotion suppression. Here, we showed that increases in delta (1-4 Hz) activity over frontal and parietal areas encompassing the brain network involved in impulse control and emotion regulation precede the occurrence of emotion suppression failures. Of note, while delta activity is considered as a typical hallmark of sleep, a growing body of evidence indicates that temporary regional increases in low-frequency, sleep-like activity may often occur also during wakefulness. Indeed, seminal work in rats demonstrated that locally synchronized neuronal *off-periods* similar to those underlying the generation of sleep slow waves (0.5-4 Hz) occur more frequently as a function of time spent awake (Vyazovskiy et al., 2011). Studies in humans also revealed that such increases are of greater magnitude in brain areas that are more intensively ‘used’ during the waking period (Hung et al., 2013; Quercia et al., 2018; Bernardi et al., 2019; Petit et al., 2019). Interestingly, the occurrence of sleep-like events in brain areas related to the execution of ongoing activities has been shown to potentially determine temporary behavioral impairments in a variety of different tasks, including impulse control, visuo-motor coordination and stimulus categorization (Bernardi et al., 2015; Nir et al., 2017). Based on these observations, it has been suggested that local sleep-like episodes may reflect the neurophysiological correlate of *‘functional fatigue’* and sleep need (Bernardi et al., 2015; Andrillon et al., 2019; D’Ambrosio et al., 2019). Our present results extend the previous literature by demonstrating that local, sleep-like episodes occurring in brain areas involved in emotion regulation predict – and thus, represent a potential cause of – failures in emotion suppression. Interestingly, our results also showed, for the first time, that sleep-like episodes related to behavioral errors do not occur only after sleep deprivation or extended task practice but may be observed also in apparently well-rested individuals, during the first hours of the morning. This finding has important implications for our understanding of the actual influence of local sleep regulation on human behavior.

### Poor sleep quality and emotion suppression failures

We found that emotion regulation failures – measured as the inability to voluntary suppress changes in facial expression -, are more common in individuals who had shorter sleep duration. Moreover, shorter sleep tended to be associated with higher delta activity over frontal regions in those who failed at suppressing their emotional expressions. These results are consistent with evidence indicating the existence of a tight interplay between sleep and emotion regulation. Indeed, both acute and chronic sleep loss determine alterations of mood and emotional reactivity, as well as increased stress, anxiety and depression (Zohar et al., 2005; Anderson and Platten, 2011; Minkel et al., 2012; Mauss et al., 2013; Beattie et al., 2015; for a recent review, also see Krause et al., 2017). The sleep-deficient individual may often display greater emotional reactivity to stimulus salience independently from valence, biased cognitive evaluation and flawed behavioral expression (Gujar et al., 2011a; Ben Simon et al., 2015). In addition, affective/mood disorders and sleep disturbances are often found associated, thus implying a potential role of inadequate sleep in the development or worsening of these clinical conditions (Benca et al., 1992; Goldstein and Walker, 2014). In line with this, good sleep quality has been associated with enhanced emotional well-being and is commonly considered as a protective factor for humans’ emotional functioning (van der Helm et al., 2011; Gruber and Cassoff, 2014; Palmer and Alfano, 2017; Watling et al., 2017).

Neuroimaging studies demonstrated that one night of sleep deprivation impairs the medial prefrontal cortex (mPFC) top-down regulation on limbic brain areas, resulting in an *‘executive dysfunction’* (Yoo et al., 2007; Gujar et al., 2011b; Gruber and Cassoff, 2014; Ben Simon et al., 2020a). Notably, even a single night of slight sleep curtailment, from 1 to 2 hours, has been shown to robustly impair mPFC activity and its related limbic functional connectivity and to determine an increased emotional distress in healthy adult individuals (Killgore, 2013). In light of these considerations, our results indicate that alterations in the top-down control exerted by the mPFC on limbic areas may be ultimately determined by the occurrence of local sleep-like episodes in this brain region.

### Limitations

It is important to acknowledge that this study has some limitations. First, while the original sample included 19 subjects, some of the analyses have been performed on a reduced sample of 12 subjects. This depended on the fact that emotion suppression failures are relatively uncommon in healthy young and rested individuals. Importantly, however, the same analyses repeated after including additional experimental sessions and subjects (N=15) confirmed the results obtained in the reduced sample (Figure S3). In future investigations, sleep restriction or deprivation paradigms could be used to increase the incidence of emotion suppression failures and thus obtain a greater statistical power.

Moreover, here we only used emotional stimuli with a positive valence, thus limiting the possibility to generalize our results to other emotional stimuli, such as negative and/or aversive stimuli. However, it should be noted that the expressive suppression strategy has been shown to rely on similar brain substrates for both negative and positive emotional stimuli (e.g., Hajcak and Nieuwenhuis, 2006; Dennis and Hajcak, 2009; Korb et al., 2012; Paul et al., 2013; Morawetz et al., 2017).

### Conclusions

Our study shows that emotion suppression failures are associated with temporary, sleep-like episodes occurring in key brain areas involved in emotion regulation. Importantly, given previous observations indicating that the incidence of local sleep-like episodes increases with time spent awake, the same functional mechanism may contribute to emotional dysfunctions commonly observed following sleep restriction or deprivation. Of note, while objective indices of sleep duration appeared to significantly correlate with behavioral impairment, the relationship was less clear for subjective reports of sleepiness. In line with previous findings, this observation suggests that individuals may not accurately perceive their actual level of brain functional fatigue (Benoit et al., 2018).

An impaired emotional regulation may substantially affect – and, potentially, disrupt – social interactions. In this light, understanding the mechanisms that underlie the loss of control over emotional responses may have broad, important implications. In particular, future studies should investigate whether specific individual factors may favor a faster build-up of or a greater vulnerability to local sleep-like episodes in specific brain areas or networks. Moreover, it will be important to clarify whether alterations in the local regulation of sleep need may be responsible for alterations of emotional regulation observed in psychopathological conditions.

## Supporting information

Avvenuti_ES_Supplementary_Material

## Acknowledgments

This work was supported by intramural funds from the IMT School for Advanced Studies Lucca. The authors wish to thank Andrea Leo, Monica Betta, and Giacomo Handjaras, for helpful discussion during the planning of the main research project, Francesca Setti for help with data acquisition, and Demetrio Grollero for assistance during data analysis.

