## Supplementary material for "Emotion suppression failures are associated with local increases in sleep-like activity": Avvenuti_ES_Supplementary_Material

**Figure S1**

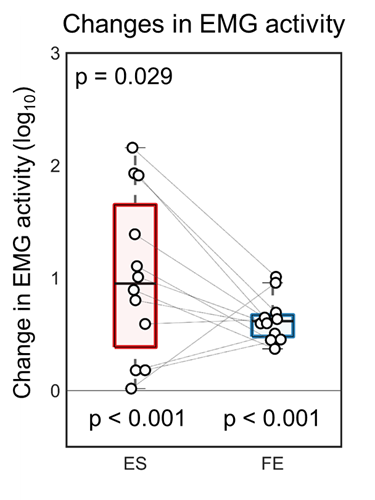

*Figure S1. Variations of EMG activity associated with changes of facial expression. Signal power changes (30-200Hz) were computed as the ratio between EMG activity in the 4 seconds after the onset of facial expression changes and brain activity in the 4 seconds that preceded this event. Values were log-transformed for display purposes. Here, values greater than 0 indicate an increase.*

**Figure S2**

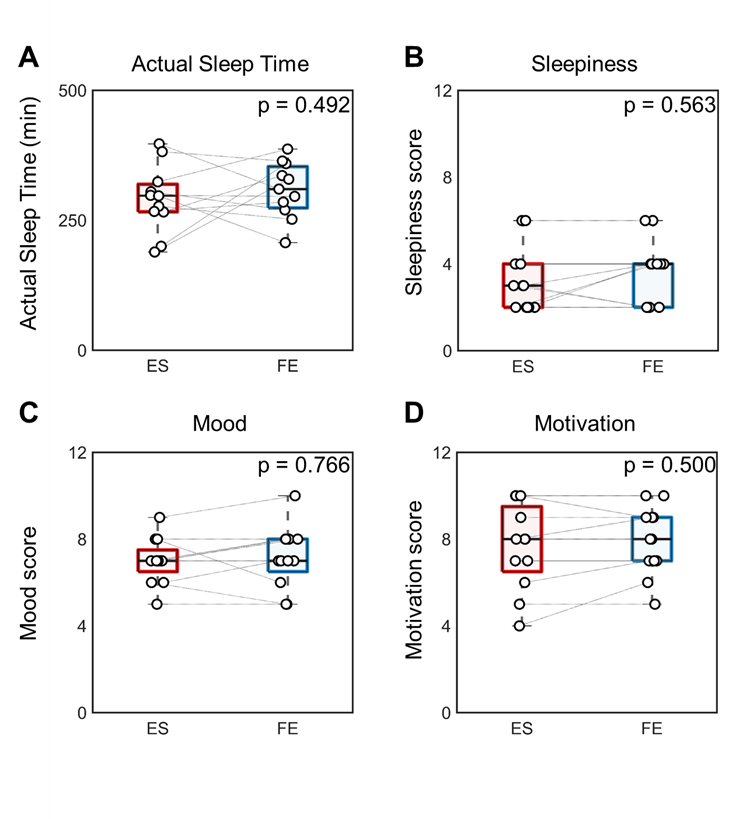

*Figure S2. (A) Total minutes of sleep in the night that preceded each experimental session (N=11). (B-D) Subjective scores (0-10) for sleepiness, mood and motivation in the two experimental conditions (N=12). No systematic differences were found between any of the tested parameters.*

**Figure S3**

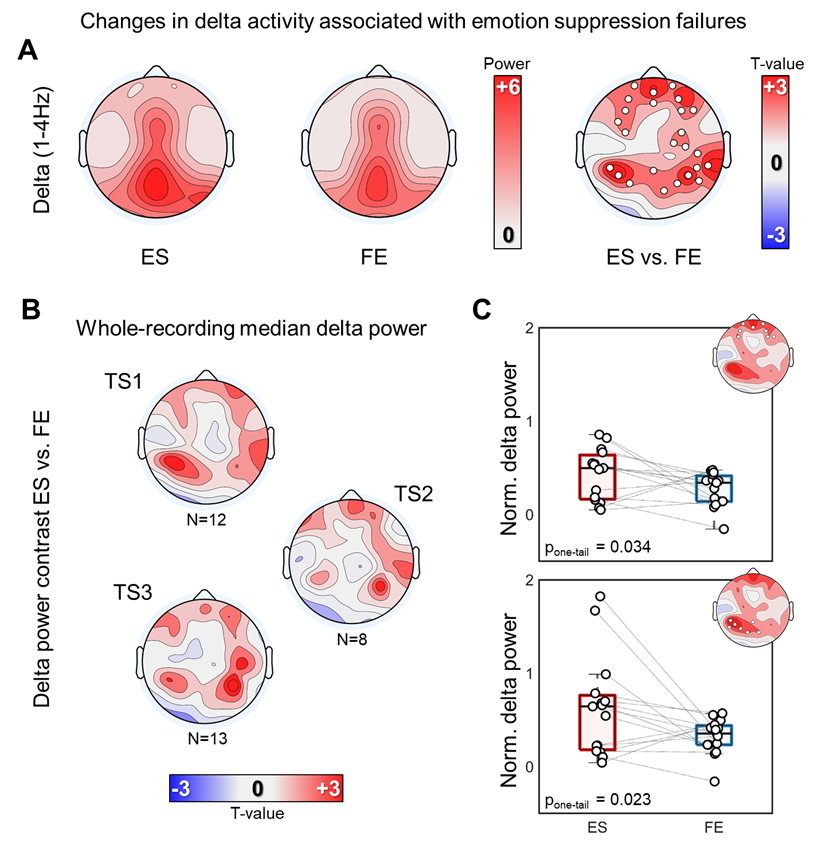

*Figure S3. (A) Delta EEG activity associated with emotion suppression failures. Topographic plots on the left show values of delta activity computed for the two experimental conditions (ES, FE) in the 4 seconds that preceded a change in facial expression. Here 15 subjects that presented at least one emotion suppression failure across the three task sessions of ES were included. (B-C) In order to take into account potential inter-session variability, additional analyses were performed after a normalization of the values of each electrode achieved by subtracting the median of delta activity computed in the whole corresponding EEG recording. The median was preferred over the mean as this measure is less likely to be affected by potential outliers. As shown in (B), median delta activity was not significantly different across experimental conditions for TS1 (12 subjects), TS2 (8 subjects) and TS3 (13 subjects). Boxplots in (C) represent differences in mean delta activity computed for the frontal and the temporo-parietal clusters obtained from the comparison of delta activity in ES and FE for the twelve subjects who presented at least one emotion suppression failure in TS1 (see main text). In topographic analyses, white dots mark p < 0.05, cluster-based correction.*

**Figure S4**

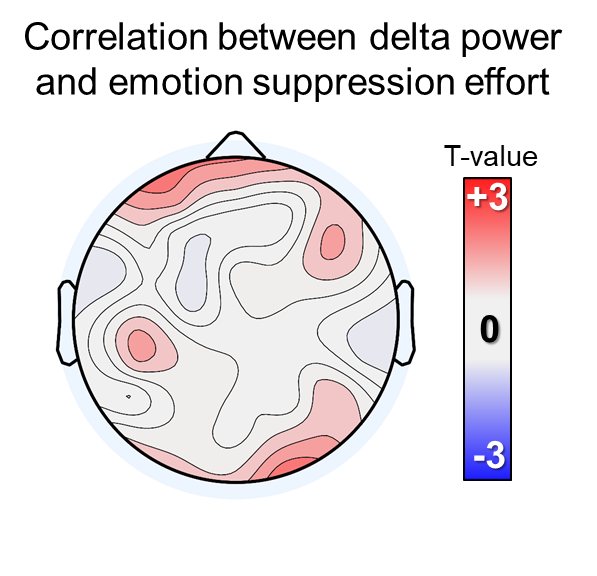

*Correlation between delta power and emotion suppression effort. For each subject, we calculated the Spearman’s correlation coefficients between mean signal power values during the presentation of each video-clip and the corresponding ratings expressing the effort needed to suppress emotional responses. A one-sample test based on z-scored correlation values was performed to identify potential group-level effects. This analysis was performed only on data from TS1 (N=12).*

**Figure S5**

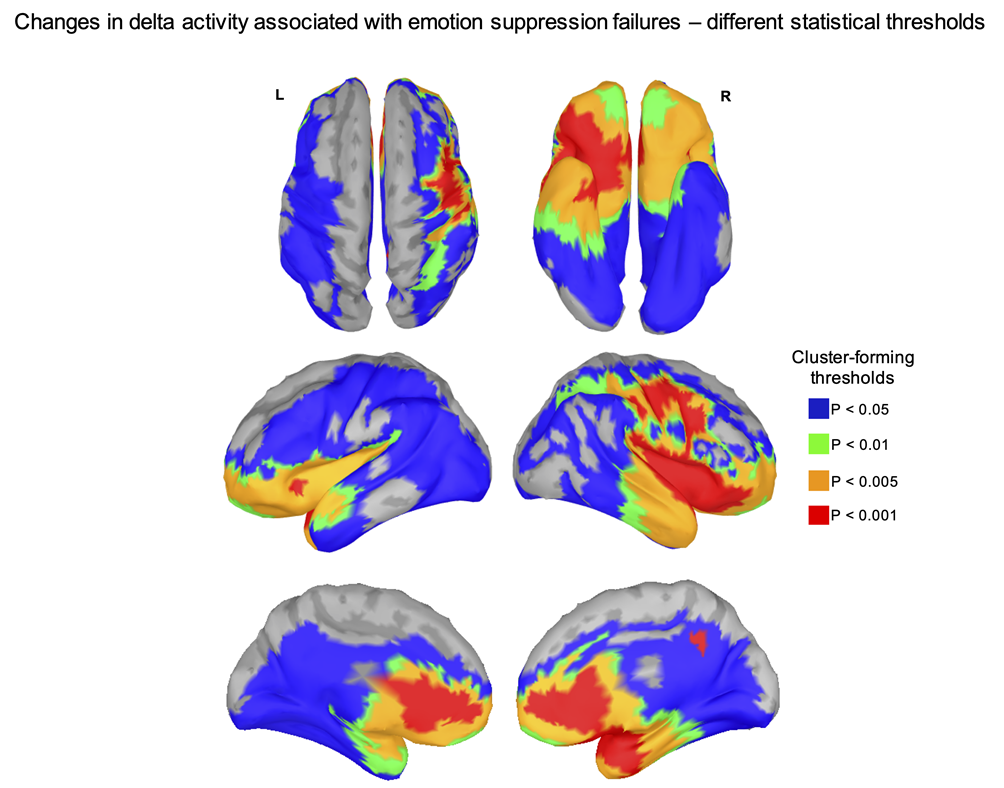

*Figure S5. Changes in delta activity associated with emotion suppression failures are reported for four different cluster-forming thresholds, corresponding to p < 0.05 (blue), p < 0.01 (green), p < 0.005 (orange), p < 0.001 (red). The cluster-level significance threshold was p < 0.05 in all cases.*
